# Ad26.COV2.S Prevents SARS-CoV-2 Induced Pathways of Inflammation and Thrombosis in Hamsters

**DOI:** 10.1101/2021.09.30.462514

**Authors:** Malika Aid, Samuel J. Vidal, Cesar Piedra-Mora, Sarah Ducat, Chi N. Chan, Stephen Bondoc, Alessandro Colarusso, Carly E. Starke, Michael Nekorchuk, Kathleen Busman-Sahay, Jacob D. Estes, Amanda J. Martinot, Dan H. Barouch

## Abstract

Syrian golden hamsters exhibit features of severe disease after SARS-CoV-2 challenge and are therefore useful models of COVID-19 pathogenesis and prevention with vaccines. Recent studies have shown that SARS-CoV-2 infection stimulates type I interferon, myeloid, and inflammatory signatures similar to human disease, and that weight loss can be prevented with vaccines. However, the impact of vaccination on transcriptional programs associated with COVID-19 pathogenesis and protective adaptive immune responses is unknown. Here we show that SARS-CoV-2 challenge in hamsters stimulates antiviral, myeloid, and inflammatory programs as well as signatures of complement and thrombosis associated with human COVID-19. Notably, single dose immunization with Ad26.COV2.S, an adenovirus serotype 26 vector (Ad26)-based vaccine expressing a stabilized SARS-CoV-2 spike protein, prevents the upregulation of these pathways such that the gene expression profiles of vaccinated hamsters are comparable to uninfected animals. Finally, we show that Ad26.COV2.S vaccination induces T and B cell signatures that correlate with binding and neutralizing antibody responses. These data provide further insights into the mechanisms of Ad26.COV2.S based protection against severe COVID-19 in hamsters.

**Author Summary:** In this study, we show that vaccination with Ad26.COV2.S protected SARS-CoV-2 challenged hamsters from developing severe COVID-19 disease by attenuating excessive proinflammatory responses, such as IL-6 and IL-1, macrophages and neutrophils signaling. Ad26 vaccination also prevented the upregulation of pathways associated with thrombosis such coagulation and clotting cascades associated with infection, and the transcriptomic profiles of vaccinated animals were largely comparable to control uninfected hamsters. In contrast, SARS-CoV-2 challenged unvaccinated hamsters showed significant increase of these proinflammatory and prothrombotic pathways and significant weight loss compared to vaccinated hamsters.

## Introduction

The COVID-19 pandemic has sparked intense interest in the rapid development of vaccines as well as animal models to evaluate vaccine candidates and to define molecular and immunologic correlates of protection. We and others have reported that animal models such as rhesus macaques and hamsters can be infected with SARS-CoV-2 and show robust viral replication in the upper and lower respiratory tract, enabling studies of COVID-19 pathogenesis and prevention with vaccines (1-5).

Golden Syrian hamsters show productive viral replication, lung pathology, and mortality when challenged with SARS-CoV2 (3, 6-8), making them pertinent for vaccine evaluation. Reports of transcriptomics and proteomics profiling of blood and lung tissues from hamsters infected with SARS-CoV-2 have shown significant upregulation of interferon and proinflammatory pathways, activation of the complement system, and recruitment of neutrophils and macrophages to the lung of infected hamsters that correlates with the presence of SARS-CoV-2 viral RNA (6, 9), supporting the role of these pro-inflammatory responses in COVID-19 severity (10). Therefore, it is important to test whether vaccines developed against COVID-19 modulate the host immune and transcriptional responses and protect from excessive proinflammatory responses induced by SARS-CoV-2.

We recently demonstrated that a single immunization with Ad26.COV2.S, an adenovirus serotype 26 (Ad26) vector-based vaccine expressing a stabilized SARS-CoV-2 spike protein, elicited binding and neutralizing antibody (NAb) responses and protected hamsters against SARS-CoV-2-induced weight loss, pneumonia, and mortality (3). These preclinical data stimulated clinical trials that demonstrated immunogenicity and efficacy of Ad26.COV2.S in humans (11, 12).

In this study we performed in depth analyses of bulk RNA-Seq transcriptomic profiling of lung tissues at day 4 post SARS-CoV-2 challenge from Ad26.COV2.S vaccinated and sham unvaccinated hamsters. To characterize Ad26 vaccine-mediated protection from severe COVID-19 in hamsters, we integrated the transcriptomics data with virological data as well as adaptive immune responses elicited by Ad26 at weeks 2 and 4. We show that a single immunization with Ad26.COV2.S (also termed Ad26.S.PP) attenuated the upregulation of proinflammatory pathways and prevented the upregulation of thrombosis associated pathways such as platelet aggregation, blood coagulation and the clotting cascade. We also find that Ad26.COV2.S vaccination upregulated signatures of CD4+, CD8+, and B cell responses that correlated with the magnitude of Ad26-elicited humoral immune responses weeks following immunization. Together, these results provide new insights into the molecular and the immunological mechanisms of Ad26.COV2.S protection from severe COVID-19.

## Results

### Study design

We recently reported a study in which recombinant, replication-incompetent Ad26 vectors expressing SARS-CoV-2 Spike constructs prevented clinical disease in Syrian golden hamsters after challenge (3). We studied two SARS-CoV-2 Spike constructs administered at either 10^10^ or 10^9^ viral particles (vp) as well as a sham control (5 total groups, n=10 per group). One Spike construct encoded a deletion of the transmembrane region and cytoplasmic tail reflecting the soluble ectodomain with a foldon trimerization domain (S.dTM.PP), while the other encoded the full-length spike (S.PP; renamed Ad26.COV2.S for clinical development). Both constructs contained two stabilizing proline mutations at the furin cleavage site. All 50 hamsters were challenged at four weeks with 5.0 × 10^5^ TCID50 of the USA-WA1/2020 strain. While both constructs prevented severe weight loss, we observed that the S.PP construct exhibited superior immunogenicity and protection (3). To gain insights into the molecular mechanisms of severe COVID-19 and Ad26-mediated protection in Syrian golden hamsters, we euthanized a subset of these animals at 4 days post-infection (4 dpi) with SARS-CoV-2 and performed bulk RNA sequencing (RNA-seq) from lung tissues. Specifically, we studied three sham unvaccinated hamsters, four S.dTM.PP-vaccinated hamsters (two at 10^10^ and two at 10^9^), and five S.PP-vaccinated hamsters (two at 10^10^ and three at 10^9^). For comparison, we additionally obtained lung tissues from three naïve animals that were neither vaccinated with Ad26 nor challenged with SARS-CoV-2 (**S1A Fig**).

### Ad26 vaccination abolishes detection of SARS-CoV-2 reads by bulk RNA-Seq

Transcriptomic profiling using bulk RNA-Seq analysis showed that the transcriptomic profiles of animals within each group of hamsters were homogenous, with no significant differences between groups of animals that received different vaccine doses (**S1B and S1C Fig**). We mapped the RNA-Seq reads to the SARS-CoV-2 genome and observed a significant number of reads that mapped to SARS-CoV-2 transcripts, ranging from 300 to 315000 reads, in the sham unvaccinated hamsters. However, we observed lower number of reads were mapped to SARS-CoV-2 transcripts in the S.dTM.PP-vaccinated hamsters, ranging from 10 to 16000 reads (**Fig 1A and S1D Fig**). In contrast, SARS-CoV-2 reads were undetectable in the Ad26.S.PP vaccinated hamsters with only 0-12 reads mapping SARS-CoV-2 transcripts (**Fig 1A and S1D Fig**). Consistent with these observations, we previously showed that detection of SARS-CoV-2 Envelope gene subgenomic RNA (sgRNA) by quantitative polymerase chain reaction (qPCR) was markedly diminished by Ad26 vaccination (3), validating our RNA-seq data. Differential expression genes analysis (DEGs) showed differences in genes upregulated or downregulated in vaccinated and sham unvaccinated animals compared to naïve animals (**S2A Fig**). Moreover, we observed significant differences in the transcriptomic profile of hamsters vaccinated with the Ad26.S.PP compared to the sham unvaccinated animals with 3401 differentially expressed genes between the two groups, whereas, only 87 genes were differentially expressed between the Ad26.S.dTM.PP vaccinated hamsters compared to sham-unvaccinated animals (**S2B Fig**) (Wald test, BH-adjust P<0.05). Moreover, we observed downregulation of proinflammatory markers such as Mx2, Irf3, Tlr7, Cxcl10 and Cd68 in the Ad26.S.PP compared to the Ad26.S.dTM.PP vaccinated hamsters (**S2C Fig)**. The expression of interferon receptors, Ifnar1 and Ifnar2 and interferon Regulatory Factor 2-Binding Protein 2(Irf2bp2) were higher in vaccinated hamsters compared to sham unvaccinated and naïve animals (**S2D Fig)**. These observations are consistent with our previous report showing that sham hamsters and hamsters vaccinated with Ad26.S.dTM.PP experienced more weight loss, extensive pneumonia, and mortality compared to hamsters that received Ad26.S.PP vaccine (3)

**Fig 1.**
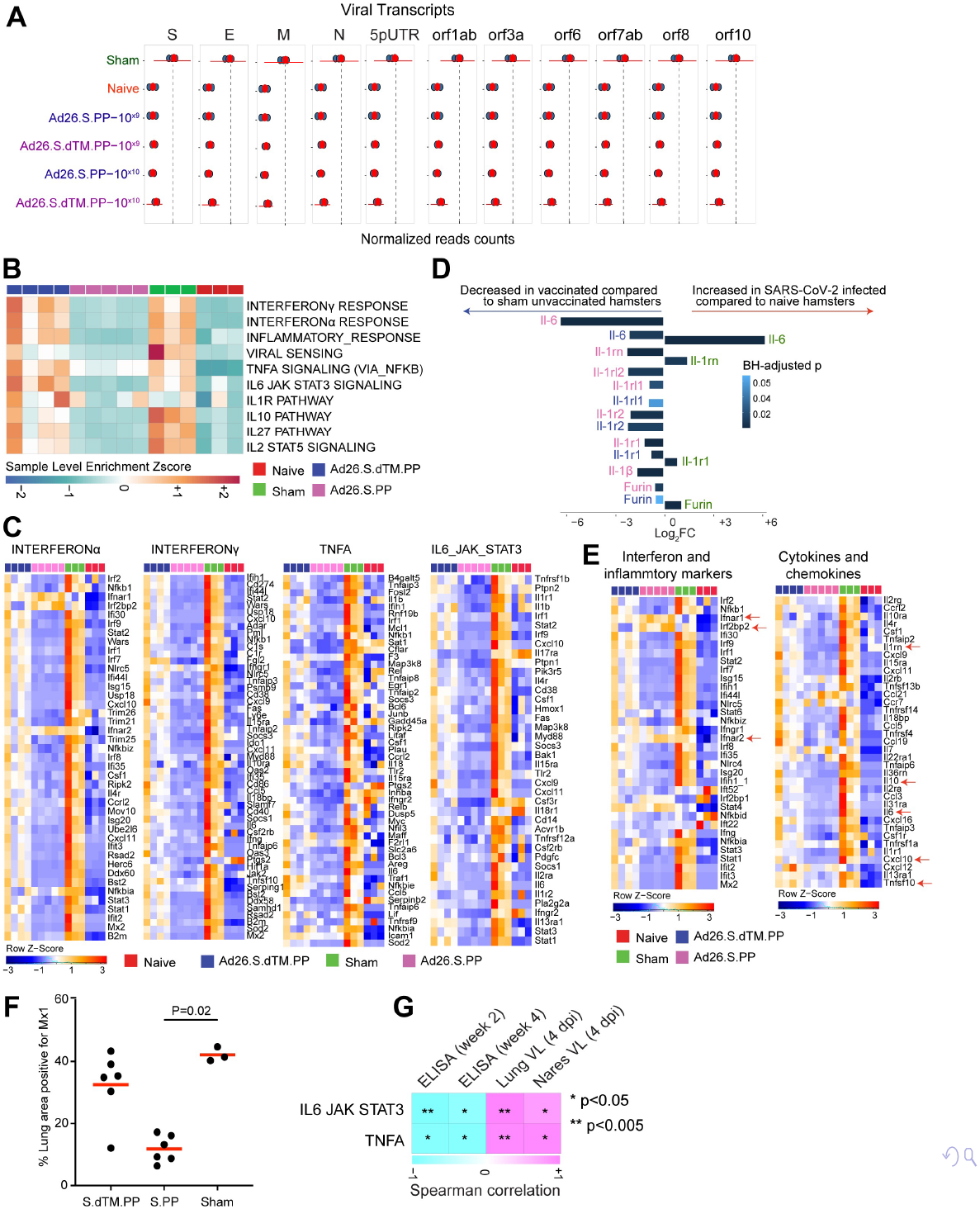
Proinflammatory signaling pathways were decreased in vaccinated compared to sham unvaccinated hamsters. **(A)** Dot plot of SARS-CoV-2 transcripts normalized read counts across all groups, vaccinated in blue and pink, sham animals in green and naïve animals in red. Each dark dot corresponds to an individual animal. Red dots represent the median normalized read counts and red horizontal bar the standard deviation for each transcript. **(B)** Sample-enrichment analysis (SLEA) of inflammatory pathways and interferon type I signaling increased by SARS-CoV-2 at 4 dpi (in green) and decreased in vaccinated hamsters (in pink and blue) compared to naïve uninfected animals (in red). Heatmap presenting the SLEA z-score of each of these pathways. SLEA represent the mean expression of all significant genes within each pathway for each animal. An SLEA z-score greater than 0 corresponds to a pathway for which member genes are up-regulated while an SLEA z-score inferior to 0 corresponds to a pathway with genes downregulated in that sample. Columns correspond to individual animals and rows correspond to individual pathways scaled across all animals using the z-score R function. **(C)** Heatmaps of the leading genes from pathways shown in a, increased or decreased at 4 dpi in vaccinated hamsters compared to sham and control naïve animals. Adjusted P-values (<0.05) were calculated by DEseq2 using Benjamini-Hochberg corrections of Wald test p-values. Columns correspond to individual animals and rows correspond to individual genes scaled across all animals. **(D)** Bar plots representation of the log2 fold change expression of IL-6, IL-1, IL-1R and Furin genes across all groups. The length of the bar plot represents the log^2^-fold change magnitude and the color gradient corresponds to the BH-adjusted p-values for each gene. **(E)** Heatmaps of proinflammatory markers and proinflammatory cytokines at 4 dpi in vaccinated hamsters compared to sham and control naïve animals. Adjusted P-values (<0.05) were calculated by DEseq2 using Benjamini-Hochberg corrections of Wald test p-values. Columns correspond to individual animals and rows correspond to individual genes scaled across all animals. **(F)** Percent area of lung tissue staining positive for Mx1. Mx1= myxovirus protein 1 (type 1 interferon inducible gene). **(G)** Correlation matrix plot showing the spearman correlation of interferon type I and inflammatory signatures at 4 dpi with viral loads in hamster’s lung and nares at 4 dpi, and with neutralizing and binding antibody titers at weeks 2-4 post vaccination. Negative correlations were shown in cyan and positive correlations were shown in pink. Correlations were assessed using Spearman correlation.

### Ad26 vaccination attenuates interferon and inflammatory signaling pathways following SARS-CoV-2 challenge

An excessive inflammatory response to SARS-CoV-2 is a major cause of disease severity and death in COVID-19. Pathways of type I and II interferon responses, and proinflammatory cytokines and chemokines were previously reported in blood and lung tissue of hamsters infected with SARS-CoV-2 and in COVID-19 patients (6, 13, 14) (**S3A and S3B Fig**). We interrogated our transcriptomic data in vaccinated hamsters compared to naïve and sham groups for proinflammatory pathways that have been shown to be central for the pathogenesis of severe COVID-19 (10, 15-17). Gene set enrichment analysis (GSEA) of DEGs at 4 dpi showed pathways of interferon signaling, inflammasome, and proinflammatory cytokines signaling such as interferon alpha, TNF, IL-1, and IL-6 signaling pathways, were significantly increased in sham unvaccinated compared to naïve hamsters [interferon-alpha (NES=2.74, FDR value <10^−6^); TNFA (NES=2.34, FDR <10^−6^); IL-6_JAK-STAT3 (NES=2.31, value <10^−6^); INFLAMMATORY RESPONSE (NES=2.39, FDR <10^−6^); IL1R signaling (NES=1.44, FDR=0.07)] as shown by the pathways individual score (sample level enrichment analysis: SLEA score) across all animals (**Fig 1B**). A direct comparison between vaccinated and sham unvaccinated hamsters at 4 dpi revealed significant decrease of proinflammatory pathways in the Ad26.S.PP vaccinated compared to sham unvaccinated animals [interferon-alpha (NES=-2.08, FDR<10^−6^) ;TNFA (NES=-2.47, FDR<10^−6^), IL-6_JAK-STAT3 (NES=-2.41, FDR<10^−6^); INFLAMMATORY RESPONSE (NES=-2.55, FDR<10^−6^); IL1R signaling (NES=-1.58, FDR=0.03)] (**Fig 1B and 1C**). Further, the expression of major proinflammatory cytokines and chemokines that contribute to SARS-CoV-2 pathogenesis, such as Il-6, Il-1α, Il-1β, Stat1/2/3, Furin, IL-2rg, Ccl21, Cxcl10, Csf, Tnf, Ccr6 and Ccr7, were significantly increased in sham unvaccinated (shown in green) compared to naïve hamsters (Wald test, BH-adjust P<0.05) (**Fig 1D and 1E)**. Whereas, the expression levels of these proinflammatory markers is comparable between vaccinated and naïve animals (**Fig 1C and 1E**).

Reduction of interferon signaling in vaccinated hamsters was validated by immunohistochemistry (IHC) for the interferon-inducible protein Mx1 in the lung of Ad26.S.PP vaccinated hamsters compared to sham infected unvaccinated hamsters (p=0.02) (**Fig 1F)**. Furthermore, we observed that proinflammatory pathways of IL6 JAK-STAT3 and TNF alpha signaling correlated negatively with Ad26 elicited ELISA binding and NAbs at weeks 2-4 following immunization (**Fig 1G and S4 Fig)**. In contrast, these pathways correlated positively with viral loads in the lung and nares of vaccinated hamsters (**Fig 1G, S4 Fig**).

### Ad26 vaccination attenuates upregulation of pathways of macrophage activation, monocytes and neutrophils

Previous bulk and single cell transcriptomics and proteomics studies in COVID-19 patients, macaques and hamsters infected with SARS-CoV-2 showed upregulation of macrophage and neutrophils signature that correlate with disease severity (4, 6, 10, 14, 16-18). We observed that pathways related to monocyte (NES=1.79, FDR=0.006), macrophage M1(NES=2.41, FDR<10^−6^) and M2 (NES=1.86, FDR=0.008), and neutrophil (NES=1.58, FDR=0.04) signaling were significantly increased in sham unvaccinated hamsters compared to naïve animals (**Fig 2A**). Vaccination with Ad26 prevented the upregulation of these pathways in the Ad26.S.PP group at 4 dpi (**Fig 2A and 2B**), whereas these pathways were slighter upregulated in the Ad26.S.dTM.PP vaccinated group (**Fig 2A and 2B**).

**Fig. 2.**
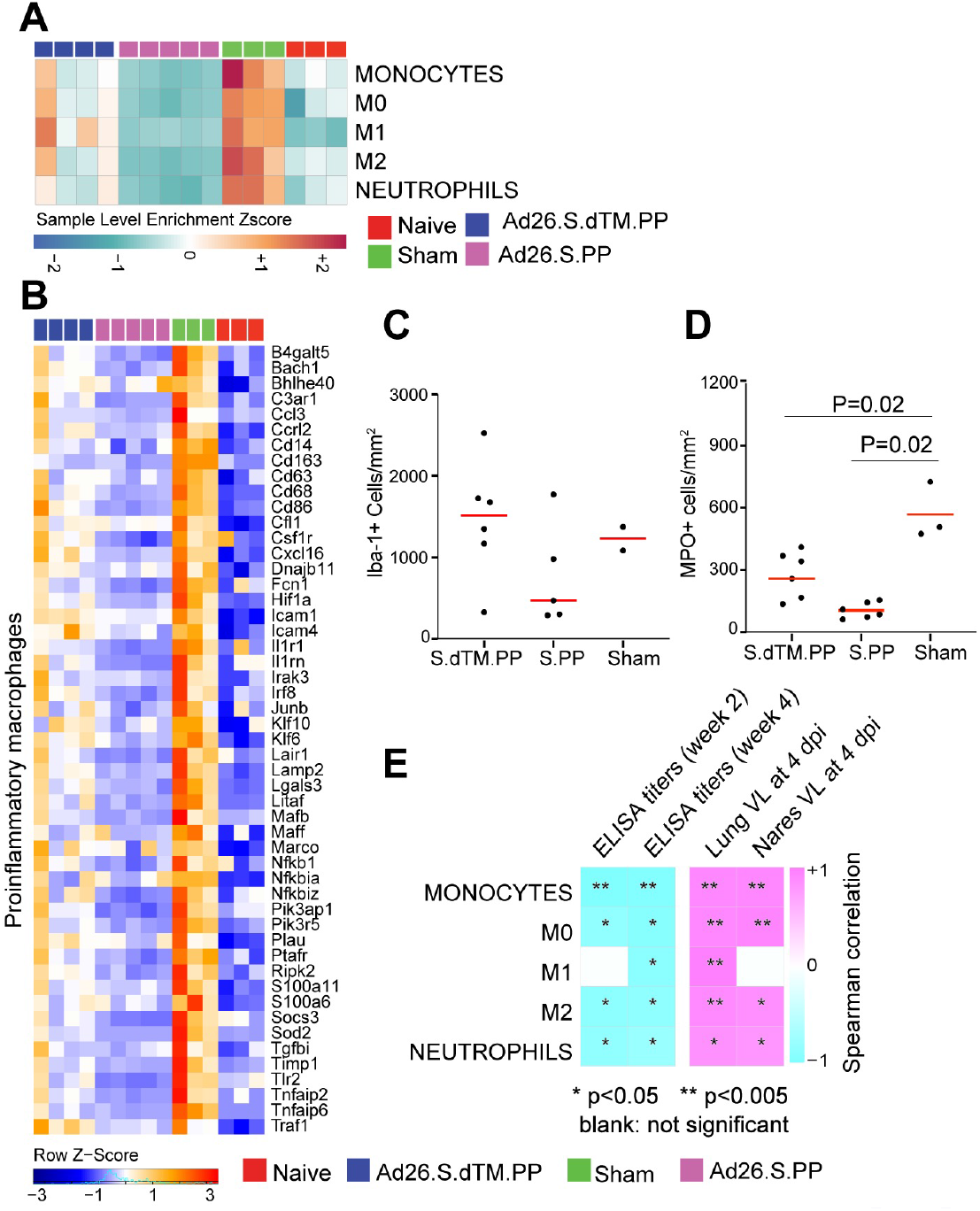
Pathways of macrophages, and neutrophils were decreased in Ad26 vaccinated compared to sham unvaccinated hamsters. **(A)** Sample-enrichment analysis (SLEA) of pathways of monocytes, macrophages M1 and M2 pathways and neutrophils signaling at 4 dpi in hamsters infected with SARS-CoV-2. In green, sham, naive in red and vaccinated groups in pink and blue colors. Heatmap presenting the SLEA z-score of each of these pathways. SLEA scores represent the mean expression of all significant genes within each pathway for each animal. An SLEA z-score greater than 0 corresponds to a pathway for which member genes are up-regulated while an SLEA z-score inferior to 0 corresponds to a pathway with genes downregulated in that sample. Columns correspond to individual animals and rows correspond to individual pathways scaled across all animals using the z-score R function. **(B)** Heatmaps of the normalized expression of the leading genes contributing to the enrichment of proinflammatory macrophages increased by SARS-CoV-2 in sham animals (in green) and decreased in Ad26 vaccinated hamsters (in pink and blue) at 4 dpi. **(C)** Number of Iba-1 positive cells (macrophages) per unit area. Dark dots represent individual animals. Red line represents median Iba-1 positive cells (macrophages) per unit area. **(D)** Number of MPO positive cells per unit area. MPO= myeloperoxidase. Dark dots represent individual animals. Red line represents median Iba-1 positive cells (macrophages) per unit area. **(E)** Correlation matrix plot showing the spearman correlations of monocytes, macrophages M0, M1 and M2 and neutrophils pathways at 4 dpi with viral loads in hamster’s lung and nares at 4 dpi, and with neutralizing and binding antibody titers at weeks 2-4 post vaccination in vaccinated hamsters. Negative correlations were shown in cyan and positive correlations were shown in pink. Correlations were assessed using Spearman correlation.

When compared to sham unvaccinated animals, Ad26.S.PP-vaccinated animals showed a significant down regulation of monocytes [NES=-2.36, FDR<10^−6^], macrophages M1 [NES=-2.33, FDR<10^−6^] and M2 [NES=-2.11, FDR<10^−6^] and neutrophils [NES=-2.11, FDR<10^−6^] pathways (**Fig 2A**). Further, we observed that markers of proinflammatory macrophages including Ccl3, Cd163, Cd68, Csf1r and Marco (6), were significantly increased in sham unvaccinated hamsters compared to naïve animals, while in the Ad26.S.PP vaccinated hamsters the expression of these macrophages markers was comparable to naïve animals (**Fig 2B)**. Consistent with these observations, vaccinated hamsters had fewer macrophages (Iba-1+) and neutrophils (myeloperoxidase, MPO+) by immunohistochemistry in lung, 4 days following challenge by quantitative image analysis (**Fig 2C and 2D**).

We observed that signatures of monocytes, M0, M1, M2 macrophages and neutrophils correlated negatively with Ad26 induced ELISA and NAbs titers at weeks 2-4 but were positively correlated with viral loads in the lung and nares of vaccinated hamsters (**Fig 2E and S5 Fig**).

### Ad26 vaccination attenuates signatures associated with complement activation and coagulation cascades

We next examined the effect of Ad26 vaccination on additional pathways known to play prominent roles in the pathogenesis of severe COVID-19 including activation of complement and coagulation cascades (14, 19, 20). Unvaccinated sham hamsters showed significant upregulation of pathways of clotting cascade [NES=1.93, FDR=0.001], coagulation cascade [NES=1.66, FDR=0.02], complement activation [NES=2.35, FDR<10^−6^], platelet activation and aggregation [NES=2.07, FDR=0.0002] and fibrinolysis [NES=2.07, FDR<10^−6^] compared to naïve animals as shown by the sample level enrichment analysis (SLEA) for each individual animal (**Fig 3A)**. Markers of these SARS-CoV-2 induced pathways in sham hamsters were enriched in complement components C3, C7, C2; clotting and coagulation cascade and tissue factors F3, F5, Plau, Fga; and platelet activation and aggregation markers such as Clu, Timp1, Thbs1, Sh2b2, and Vav1 (**Fig 3B)**. These markers were less or not increased in the Ad26 vaccinated hamsters (**Fig 3B)**. When compared to sham unvaccinated, vaccinated hamsters showed a significant down regulation of pathways of clotting cascade [NES=-1.98, FDR=0.001], coagulation cascade [NES=-2.23, FDR<10^−6^], complement activation [NES=-2.04, FDR=0.0007], platelet activation and aggregation [NES=-1.64, FDR=0.02] and fibrinolysis [NES=-1.5, FDR=0.05] (**Fig 3C**), suggesting Ad26 vaccination protected hamsters from developing thrombosis associated pathways. Moreover, we found that pathways of complement cascade, platelet activation and aggregation, coagulation cascade, and fibrinolysis correlated negatively with weeks 2-4 ELISA binding and neutralizing antibody titers (NAb) elicited by Ad26 vaccination, and positively correlated with 4 dpi viral loads in lung and nares of vaccinated hamsters (**Fig 3D and S6 Fig**).

**Fig. 3.**
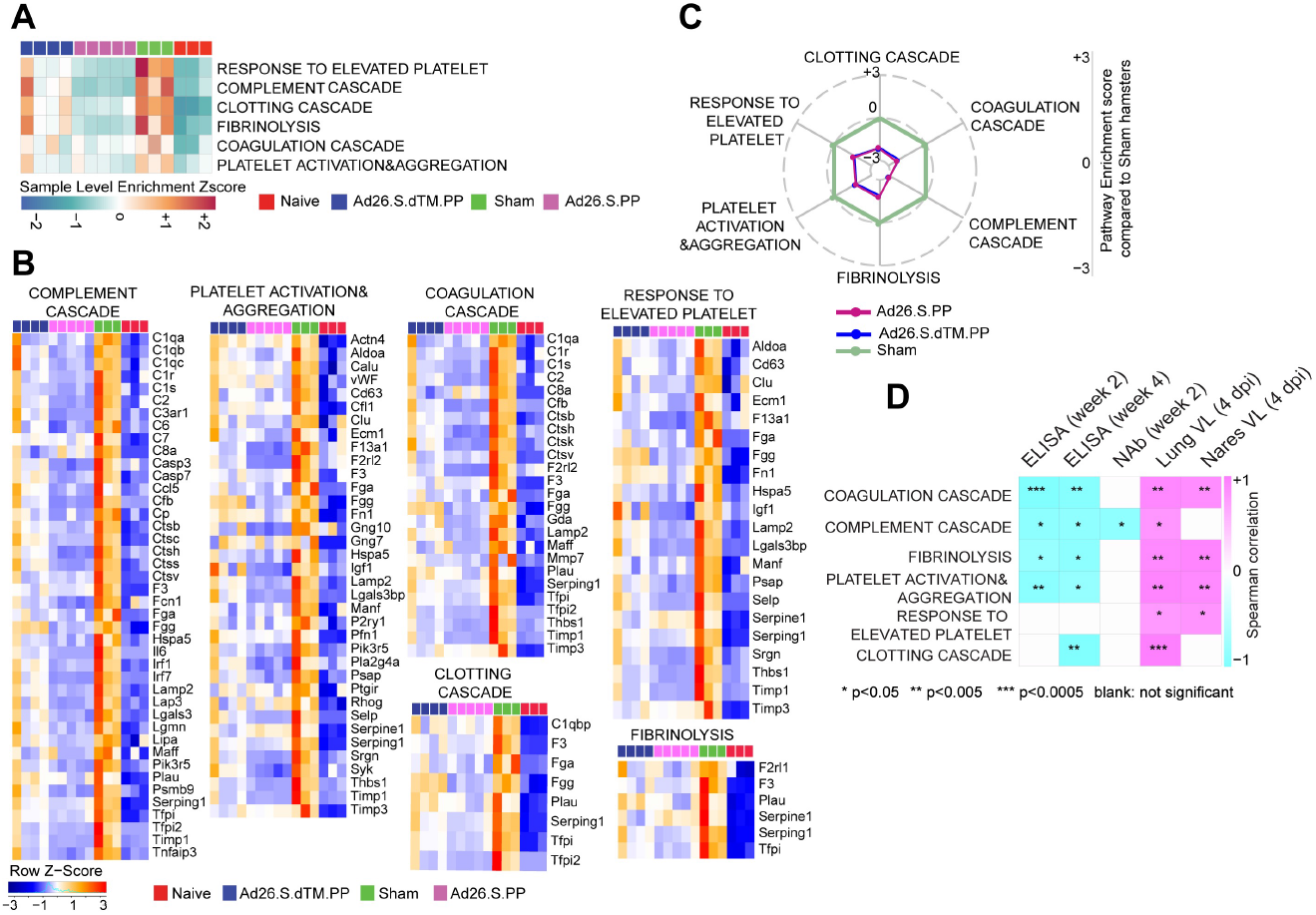
Ad26 vaccination attenuates signatures of complement system activation and coagulation cascade associated with severe COVID-19. **(A)** Sample-enrichment analysis (SLEA) of pathways of thrombosis associated pathways decreased in Ad26 vaccinated animals at 4 dpi (in pink and blue) compared to sham-unvaccinated animals (in green). Heatmap presenting the SLEA z-score of each of these pathways. SLEA represent the mean expression of all significant genes within each pathway for each animal. **(B)** Heatmaps of the normalized expression of the leading genes contributing to the enrichment of thrombosis associated pathways increased by SARS-CoV-2 in sham animals (in green) and decreased in Ad26 vaccinated hamsters (in pink and blue) at 4 dpi. **(C)** Radar plot showing the GSEA normalized enrichment score (NES) of thrombosis associated pathways decreased in Ad26 vaccinated hamsters in blue and pink compared to sham unvaccinated animals in green. NES value in the sham group were normalized to itself and were put to zero. **(D)** Correlation matrix plot showing the spearman correlations of thrombosis associated pathways at 4 dpi with viral loads in hamster’s lung and nares at 4 dpi, and with neutralizing and binding antibody titers at weeks 2-4 post vaccination. Negative correlations were shown in cyan and positive correlations were shown in pink. Correlations were assessed using Spearman correlation.

### Ad26 vaccination induces T and B cell signatures that correlated with Ad26 induced humoral immune responses

We next investigated whether Ad26 vaccination induced molecular signatures consistent with protective adaptive immune mechanisms observed in SARS-CoV-2 infected humans, macaques and hamsters vaccinated with Ad26 (3, 11, 21, 22). GSEA showed that signatures of CD4+ T cell [NES=2.27, FDR<10^−6^], CD8+ T cell [NES=1.73, FDR=0.02], T helper follicular cell (Tfh) [NES=1.83, FDR=0.009], and T regulatory cell [NES=2.07, FDR=0.0003], were enriched in Ad26-vaccinated hamsters compared to naïve controls (**Fig 4A and 4B)**. GO term analysis of CD4+ T cell pathway (Cd28, Cd3g, Cd3e, Cd3d, Lck, Fyn, Cd247, Ccr7, Dpp4, Icos), showed enrichment of markers of lymphocyte co-stimulation (FDR=2.65 × 10^−12^), leukocyte activation (FDR=7.25 x10^−12^), and T cell activation (FDR=1.18 × 10^−10^). Moreover, key markers of cytotoxic CD8+ T cell adhesion, activation, and effector memory functions such as Prf1, Lef1, Lck, Tcf7 and Nkg7, were increased by Ad26 in vaccinated compared to naïve control animals (**Fig 4B**), suggesting CD8+ T cell-induced by Ad26 are engaged in the recognition and elimination of infected cells. Similarly, Th1 cell signatures were significantly upregulated in Ad26-vaccinated compared to naïve control animals and include Th1 cytotoxic genes (NES=2.01, FDR<10^−6^), genes increased in Th1 cells (NES=1.6, FDR=0.02) and STAT4 signaling, a major transcription factor involved in Th1 differentiation and proliferation (23) (NES=1.31, FDR=0.02) (**Fig 4C**).

**Fig. 4.**
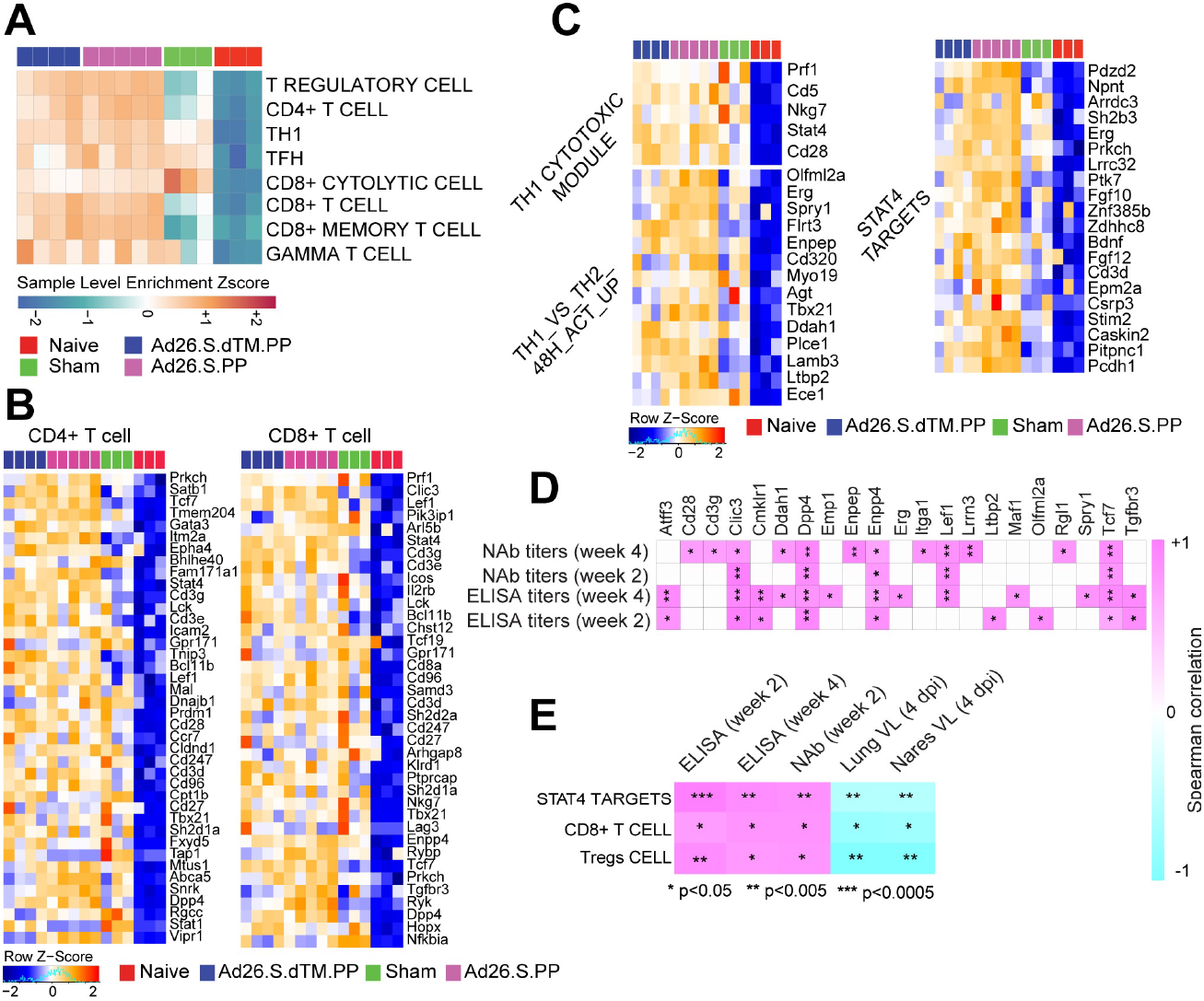
Ad26 vaccination increased pathways of CD8 and CD4 signaling in vaccinated hamsters. **(A)** Sample-enrichment analysis (SLEA) of pathways of signatures of CD4, CD8, T regulatory cell (Tregs), T follicular helper cell (TFH) and T helper 1 (Th1) cell increased in vaccinated animals at 4 dpi (in pink and blue) compared to naïve and sham animals (in red and green). Heatmap presenting the SLEA z-score of each of these pathways. SLEA represent the mean expression of all significant genes within each pathway for each animal. An SLEA z-score greater than 0 corresponds to a pathway for which member genes are up-regulated while an SLEA z-score inferior to 0 corresponds to a pathway with genes downregulated in that sample. Columns correspond to individual animals and rows correspond to individual pathways scaled across all animals using the z-score R function. **(B-C)** Heatmaps of the leading genes of CD8+, CD4+, Tregs and Th1 markers, increased at 4 dpi in vaccinated animals compared to naïve animals. Adjusted P-values (<0.05) were calculated by DEseq2 using Benjamini-Hochberg corrections of Wald test p-values. Columns correspond to individual animals and rows correspond to individual genes normalized reads count scaled across all animals. **(D-E)** Matrix showing the spearman correlations of CD4 T cell markers (D) and CD8 T cell signature, Tregs signature and STAT4 targets (E) at 4 dpi with viral loads in hamster’s lung and nares, and with neutralizing and binding antibody titers at weeks 2-4 in vaccinated hamsters. Negative correlations were shown in cyan and positive correlations were shown in pink. Correlations were assessed using Spearman correlation.

We next correlated T cell signatures induced in vaccinated hamsters with humoral immune responses elicited by the Ad26 vaccine at weeks 2-4. We observed a significant positive correlation of markers of CD4+ T cell (**Fig 4D**), signatures of CD8+ T cell and T regulatory cell signatures with ELISA and neutralizing antibody titers at weeks 2 and 4, whereas these signatures correlated negatively with viral loads in the lung and nares of vaccinated hamsters (**Fig 4E**). Similarly, we found that STAT4 targets, a major transcription factor involved in Th1 activation (23, 24), were positively correlated with ELISA and neutralizing antibody titers at weeks 2 and 4, and negatively correlated with viral loads in the lung and nares of vaccinated hamsters (**Fig 4E**).

Given the critical role of humoral responses in immunity to SARS-CoV-2 infection (25), we next characterized B cell responses following Ad26 vaccination in hamsters. We observed a significant increase in signatures of B cell activation and differentiation and other markers regulating B cell fate and development in vaccinated hamsters compared to sham and naïve animals, as shown by the SLEA scores in each individual animal (**Fig 5A**) and by the upregulation of the top markers of B cell activation and differentiation and markers regulating B cell fate and development such as Cd79, Pou2af1, Bank1and Tcf4 (**Fig 5B**). These pathways correlated positively with binding and neutralizing antibody titers elicited by Ad26 at weeks 2 and 4 (**Fig 5C and S7 Fig**) but were correlated negatively with viral loads in the lung and nares of vaccinated hamsters (**Fig 5C and S7 Fig**). Additionally, pathways of B cell activation, differentiation, and development correlated positively with S-specific and RBD-specific antibody IgG, IgG2a, IgG3, IgM, Fc-receptors FcRγ2, FcRγ3, and FcRγ4 and antibody-dependent complement deposition (ADCD) responses in vaccinated animals at week 4 assessed by systems serology and previously published by our group (3) (**Fig 5D**).

**Fig. 5.**
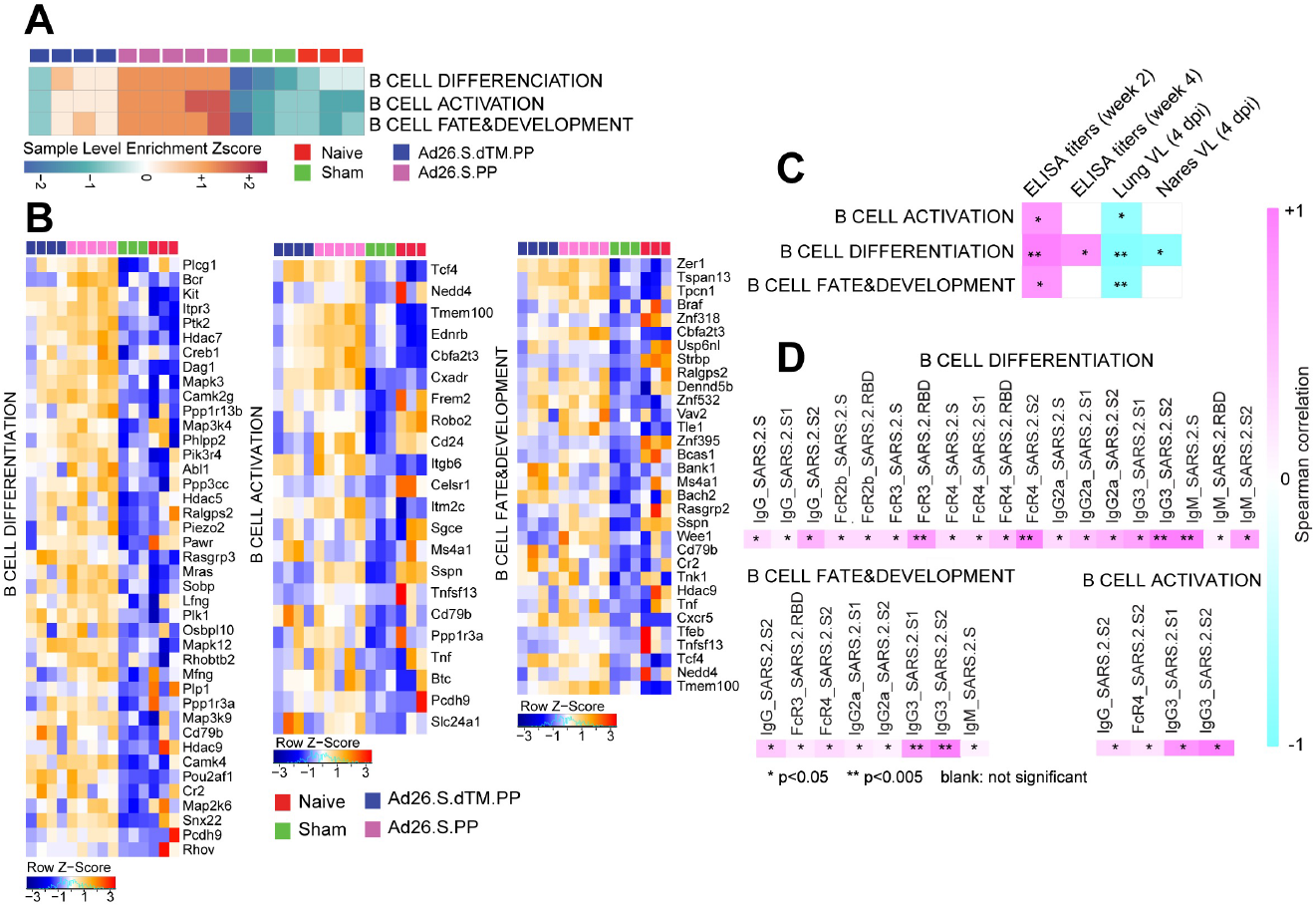
Pathways of B cell increased in vaccinated compared to naïve and sham hamsters. **(A)** Sample-enrichment analysis (SLEA) of pathways of B cell activation, differentiation and B cell fate increased in vaccinated animals at 4 dpi (in pink and blue) compared to naïve and sham animals (in red and green). Heatmap presenting the SLEA z-score of each of these pathways. SLEA represent the mean expression of all significant genes within each pathway for each animal. **(B)** Heatmaps of the leading genes of B cell markers from pathways shown in the top panel, increased at 4 dpi in vaccinated compared to sham and naïve hamsters. Adjusted P-values (<0.05) were calculated by DEseq2 using Benjamini-Hochberg corrections of Wald test p-values. Columns correspond to individual animals and rows correspond to individual gene normalized reads count scaled across all animals. **(C)** Matrix showing the spearman correlation of B cell signatures at 4 dpi with viral loads in hamster’s lung and nares, and with neutralizing and binding antibody titers at weeks 2-4 in vaccinated hamsters. Negative correlations were shown in cyan and positive correlations were shown in pink. Correlations were assessed using Spearman correlation. **(D)** Matrix showing the Spearman correlations of B cell pathways with S-specific and RBD-specific antibody responses in the Ad26.S.PP vaccinated animals at week 4 by systems serology, including IgG, IgG2a, IgG3, IgM, Fc-receptors FcRγ2, FcRγ3, FcRγ4, and antibody-dependent complement deposition (ADCD). All positive correlations were shown in pink. Correlations were assessed using Spearman correlation.

## Discussion

We previously showed that vaccination with Ad26.COV2.S, an Ad26 vector encoding a full-length prefusion stabilized S immunogen (S.PP), protected against SARS-CoV-2 challenge in hamsters (3) and macaques (21). Here we used bulk RNA-Seq of the lung, and showed that Ad26-based vaccines both attenuate hallmark pathologic pathways associated with COVID-19 and stimulate signatures of protective adaptive immune responses. These observations are consistent with a decrease of lung injury and tissue damage in vaccinated hamsters reported previously (3) (26) and highlight the protective role of Ad26 vaccination from SARS-CoV-2 pathology in both macaques and hamsters (3, 21).

In a Phase 1/2a randomized, clinical study, Ad26.COV2.S vaccination generated robust CD8+ T cell, CD4+ T cell, and antibody responses following vaccination (11, 12). Consistent with these clinical data, in the Ad26 vaccinated hamsters we observed upregulation of signatures of effector and cytolytic CD8+ T cell, CD4+ T cell signatures. More importantly, we showed that these signatures correlated positively with binding and neutralizing antibody titers elicited by Ad26 at weeks 2 and 4 and negatively with viral loads in lungs and nares at 4 dpi.

In line with the Ad26 vaccination protective role against SARS-CoV-2 infection, we observed a pronounced increase of signatures of B cell activation, differentiation, and development in vaccinated compared to sham and naïve hamsters at 4 dpi, with slightly higher expression levels in Ad26.S.PP vaccinated hamsters compared to hamsters vaccinated with Ad26.S.dTM.PP. We showed that B cell signatures induced by Ad26 correlated positively with neutralizing and binding antibody titers and with S-specific and RBD-specific antibody IgG, IgG2a, IgG3, IgM, Fc-receptors FcRγ2, FcRγ3, and FcRγ4 and antibody-dependent complement deposition (ADCD) responses at week 4 following vaccination. This indicates successful humoral effector programs induced by the Ad26.S.PP vaccine that correlated with protection against SARS-CoV-2 infection (25).

Overall, vaccination with Ad26.COV2.S prevented the upregulation of the pathological pathways induced by SARS-CoV-2, and the transcriptomic profile of vaccinated animals was largely comparable to sham uninfected hamsters. Ad26 immunization in hamsters also led to the induction of cellular and humoral immune responses that provide insights into the potential mechanisms of protection in lungs against SARS-CoV-2.

## Materials and Methods

### Animals and study design

Seventy male and female Syrian golden hamsters (Envigo), 10–12 weeks old, were randomly allocated to groups. All animals were housed at Bioqual. Animals received Ad26 vectors expressing S.dTM.PP or S.PP or sham controls (n = 10 per group). Animals received a single immunization of 10^10^ or 10^9^ vp Ad26 vectors by the intramuscular route without adjuvant at week 0. At week 4, all animals were challenged with 5.0 × 10^5^ TCID50 or 5.0 × 10^4^ TCID50 SARS-CoV-2, which was derived with one passage from USA-WA1/2020 (NR-52281, BEI Resources). Virus was administered as 100 μl by the intranasal route (50 μl in each nare). Body weights were assessed daily. All immunologic and virologic assays were performed blinded. All animal studies were conducted in compliance with all relevant local, state and federal regulations and were approved by the Bioqual Institutional Animal Care and Use Committee. On day 4, a subset of animals was euthanized for tissue viral loads, pathology and transcriptomics profiling. We performed bulk RNA-Seq on lung frozen tissues from 3 naïve controls, 3 SARS-CoV-2 infected animals, 4 animals vaccinated with Ad26 vectors expressing S.dTM.PP and 5 animals vaccinated with Ad26 vectors expressing S.PP. The immunological, virological and humoral data were previously published by our group (3).

Lung tissue was homogenized in 700 μL of QIAzol (Qiagen) and stored at -80°C until being extracted using the miRNeasy Micro kit (Qiagen) with on-column DNase digestion. RNA quality was assessed using an Agilent Bioanalyzer and ten nanograms of total RNA used as input for library preparation using the SMARTer Stranded Total RNA-Seq V2 Pico Input Mammalian kit (Takara Bio) according to the manufacturer’s instructions. Libraries were validated by capillary electrophoresis on an Agilent 4200 TapeStation, pooled at equimolar concentrations, and sequenced on an Illumina NovaSeq6000 at 100SR, targeting 25-30 million reads per sample. Alignment was performed using STAR version 2.7.3a (27) with the MesAur1.0 (GCF_000349665.1) assembly and annotation of the hamster downloaded from NCBI. Transcript abundance estimates was calculated internal to the STAR aligner using the algorithm of htseq-count as described previously (4). DESeq2 was used for normalization, producing both a raw and normalized read count table. Differential expression at the gene level were performed by DESeq2 implemented in the DESeq2 R package. A corrected p-value cut-off of 0.05 was used to assess significant genes that were upregulated or down regulated by SARS-Cov2 at day 4 post challenge in sham and vaccinated animals compared to naïve controls and in vaccinated compared to sham animals using Benjamini-Hochberg (BH) method.

### Histopathology and immunohistochemistry

Tissues were fixed in freshly prepared 4% paraformaldehyde for 24 hours, transferred to 70% ethanol, paraffin embedded within 7-10 days, and blocks sectioned at 5 μm. Slides were baked for 30-60 min at 65°C then deparaffinized in xylene and rehydrated through a series of graded ethanol to distilled water. For Iba-1 IHC, heat induced epitope retrieval (HIER) was performed using a pressure cooker on steam setting for 25 minutes in citrate buffer (Thermo; AP-9003-500) followed by treatment with 3% hydrogen peroxide. Slides were then rinsed in distilled water and protein blocked (BioCare, BE965H) for 15 min followed by rinses in 1x phosphate buffered saline. Primary rabbit anti-Iba-1 antibody (Wako; 019-19741 at 1:500) was applied for 30 minutes followed by rabbit Mach-2 HRP-Polymer (BioCare; RHRP520L) for 30 minutes then counterstained with hematoxylin followed by bluing using 0.25% ammonia water. Labeling for Iba-1 was performed on a Biogenex i6000 Autostainer (v3.02). In some cases, Iba-1 staining was performed with Iba-1 at 1:500 (BioCare Cat. No. CP290A; polyclonal), both detected by using Rabbit Polink-1 HRP (GBI Labs Cat. No. D13-110). Neutrophil (MPO), and type 1 IFN response (Mx1) was performed with MPO (Dako Cat. No. A0398; polyclonal) at 1:1000 detection using Rabbit Polink-1 HRP, and Mx1 (EMD Millipore Cat. No. MABF938; clone M143/CL143) at 1:1000 detection using Mouse Polink-2 HRP (GBI Labs Cat. No. D37-110). Staining for MPO and Mx1 IHC was performed as previously described using a Biocare intelliPATH autostainer, with all antibodies being incubated for 1 h at room temperature.

Quantitative image analysis was performed using HALO software (v2.3.2089.27 or v3.0.311.405; Indica Labs) on at least one lung lobe cross section from each animal. In cases where >1 cross-section was available; each lung lobe was quantified as an individual data point. For Mx1 quantification, the Area Quantification v2.1.3 module was used to determine the percentage of Mx1 protein as a proportion of the total tissue area. For Iba-1 quantification, the Multiplex IHC v2.3.4 module was used to detect Iba-1+ cells and is presented as a proportion of total alveolar tissue. For MPO (neutrophil) quantification, the HALO AI software was first trained to detect the alveolar portion of the lung by excluding blood vessels (>5mm^2^), bronchi, bronchioles, cartilage, and connective tissue; subsequently, the Multiplex IHC v2.3.4 module was used to detect MPO+ cells and is presented as a proportion of total alveolar tissue (PMNs/mm^2^). In all instances, manual curation was performed on each sample to ensure the annotations were accurate and to correct false positives/false negatives.

### Pathway enrichment analyses

Gene set enrichment analysis and a compendium of databases of biological and immunological pathways were used to test the longitudinal enrichment of pathways and transcription factors (TFs) signatures at 4 dpi in vaccinated, sham and control hamster groups. Genes were pre-ranked by fold change from the highest to the lowest and GSEA was used to assess the enrichment of selected gene sets. Thrombosis signatures were compiled from the MSigDB curated C2 gene sets, IPA ingenuity pathway analysis (https://targetexplorer.ingenuity.com) and were manually curated in-house by checking the individual function of each gene using GeneCard data base (https://www.genecards.org/). Cytokines signaling, immune cell signatures, and molecular pathways were compiled from the MSigDB Hallmark, C2, C7 and C3 gene sets (https://www.gsea-msigdb.org/gsea/msigdb/collections.jsp), IPA ingenuity pathway analysis https://targetexplorer.ingenuity.com), and the blood transcriptional modules (BTMs)(28). The GSEA Java desktop program was downloaded from the Broad Institute (http://www.broadinstitute.org/gsea/index.jsp) and used with GSEA Pre-Ranked module parameters (number of permutations: 1,000; enrichment statistic: weighted; 10≤ gene set size ≤5,000). Sample-level enrichment analysis SLEA (29) was used to investigate the enrichment of pathways in each animal. Briefly, the expression of all the genes in a specific pathway was averaged across samples and compared to the average expression of 1,000 randomly generated gene sets of the same size. The resulting z-score was then used to reflect the overall perturbation of each pathway in each sample.

### Canonical pathway and upstream regulator functions analysis

The canonical pathway and upstream regulator functions of IPA core expression analysis tool (Qiagen) were used to interrogate the lists of genes upregulated or down regulated by SARS-CoV-2 at 4 dpi in sham infected hamsters compared to macaques infected with SARS-CoV-2. Canonical pathways and upstream regulators were considered significant if pathway activation Z-Score ≥ 2 and pathway overlap corrected p-value < 0.05 (using the Benjamini-Hochberg method). Functional analysis of statistically significant gene and protein changes was performed using Ingenuity Pathways Knowledge Base (IPA; Ingenuity Systems). For all gene set enrichment analyses, a right-tailed Fisher’s exact test was used to calculate P-values associated with each biological function and canonical pathway. The calculated z-score signifies whether a gene or protein expression changes for known targets of each regulator are consistent with what is expected from the literature (z > 2, regulator predicted to be activated, z < -2, regulator predicted to inhibited). Additional functional module analyses were performed using Functional module detection (https://hb.flatironinstitute.org/module/). Go term enrichment analysis was performed using the GeneMania database (https://genemania.org/). For GSEA analysis, all significant pathways and molecular signatures, up-or down-regulated in the different groups of hamsters, were selected using a false discovery rate < 20 and a nominal p-value < 0.05. In IPA Global Canonical Pathways (GCP), a multiple-testing corrected p-value was calculated using the Benjamini-Hochberg (BH) method.

### Quantification and statistical analysis

Quality of RNA-Seq raw reads was examined using FastQC (http://www.bioinformatics.babraham.ac.uk/projects/fastqc/) and reads were aligned using STAR v2.7.3. Differential expression at the gene level was performed by DESeq2 implemented in the DESeq2 R package. Pathways enrichment and upstream regulators analyses were analyzed through the use of ingenuity pathway analysis (IPA) (QIAGEN Inc., https://www.qiagenbioinformatics.com/products/ingenuitypathway-analysis) and the GSEA Desktop v4.0.3 from (https://www.gsea-msigdb.org/gsea/index.jsp). For data annotation and presentation, we used a collection of tools including Cytoscape version 3.6.0 (https://cytoscape.org), GeneMANIA version 3.3.1 (http://genemania.org) and Genecards (https://www.genecards.org). Where indicated on the figures, all p-values were adjusted for multiple comparisons using the Benjamini-Hochberg method (BH). Correlations of transcriptomics signatures with viral loads, ELISA binding, and neutralizing titers were measured by correlating the pathway sample level enrichment score (SLEA) with the different outcomes using two-sided Spearman_s rank correlations. All statistical analyses were performed using the R statistical software 3.5.1. All analyses were conducted using R programming language.

## Acknowledgments

We thank Kathryn L. Pellegrini, Gregory Tharp, Steven E. Bosinger, Hanneke Schuitemaker, Frank Wegmann and Roland Zahn for generous advice, assistance, and reagents.

We acknowledge support from the Ragon Institute of MGH, MIT, and Harvard, Mark and Lisa Schwartz Foundation, Beth Israel Deaconess Medical Center, Massachusetts Consortium on Pathogen Readiness (MassCPR), Bill & Melinda Gates Foundation (INV-006131), and the Janssen Vaccines & Prevention program.

## Supporting information captions

**S1 Fig.**
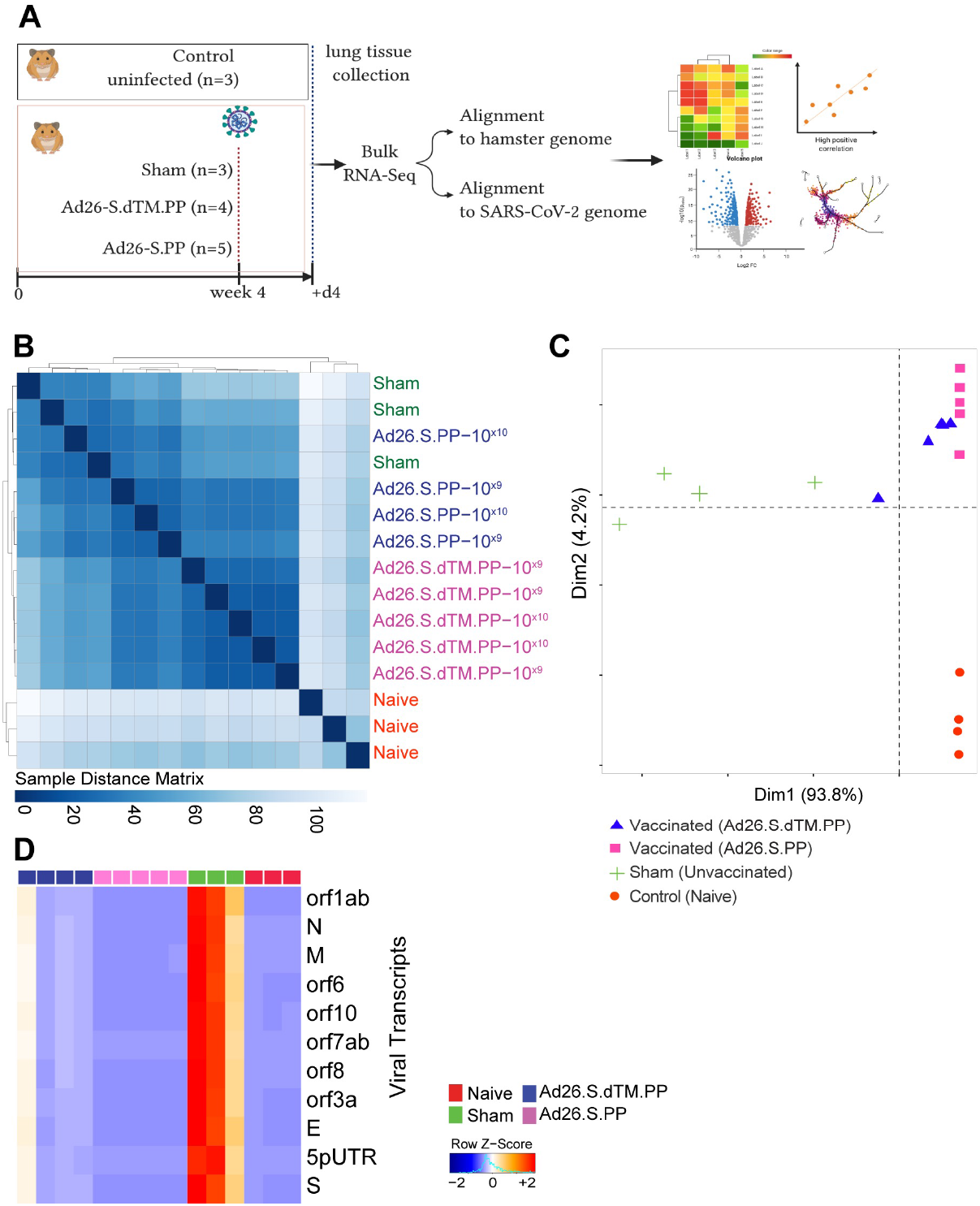
Transcriptomic profiling of naïve, sham and vaccinated hamsters. **(A)** Study design. Four groups of Syrian golden hamsters were used in this study from a previously published study by our group (3). Lung snap frozen tissues were sampled at day 4 post challenge for bulk RNA-Seq. **(B)** Expression similarity matrix comparing all animals across the four groups. Hierarchical clustering showing uninfected control animals in red in a separate cluster from sham unvaccinated and vaccinated animals. The distance matrix was generated using the R function dist (). In pink color: animals vaccinated with the Ad26.S.PP ; in blue: animals vaccinated with Ad26.S.dTM.PP; in green: sham infected and unvaccinated animals and in red: naïve animals. **(C)** Unsupervised clustering of naïve, infected unvaccinated and vaccinated groups using PCA plot generated by the prcomp () R function on the two first components where vaccinated groups were shown in blue and pink; naïve uninfected animals in red and sham unvaccinated animals in green. **(D)** Heatmap of SARS-CoV-2 genes enriched in sham unvaccinated animals, and decreased at 4 dpi in vaccinated animals. Vaccinated groups were shown in pink and blue colors compared to sham (green) and naïve (red) animals. A blue-to-red color gradient represents the row normalized expression value for each gene across all samples. Adjusted p-values were calculated by DEseq2 using Benjamini-Hochberg (BH) corrections of Wald test p-values.

**S2 Fig.**
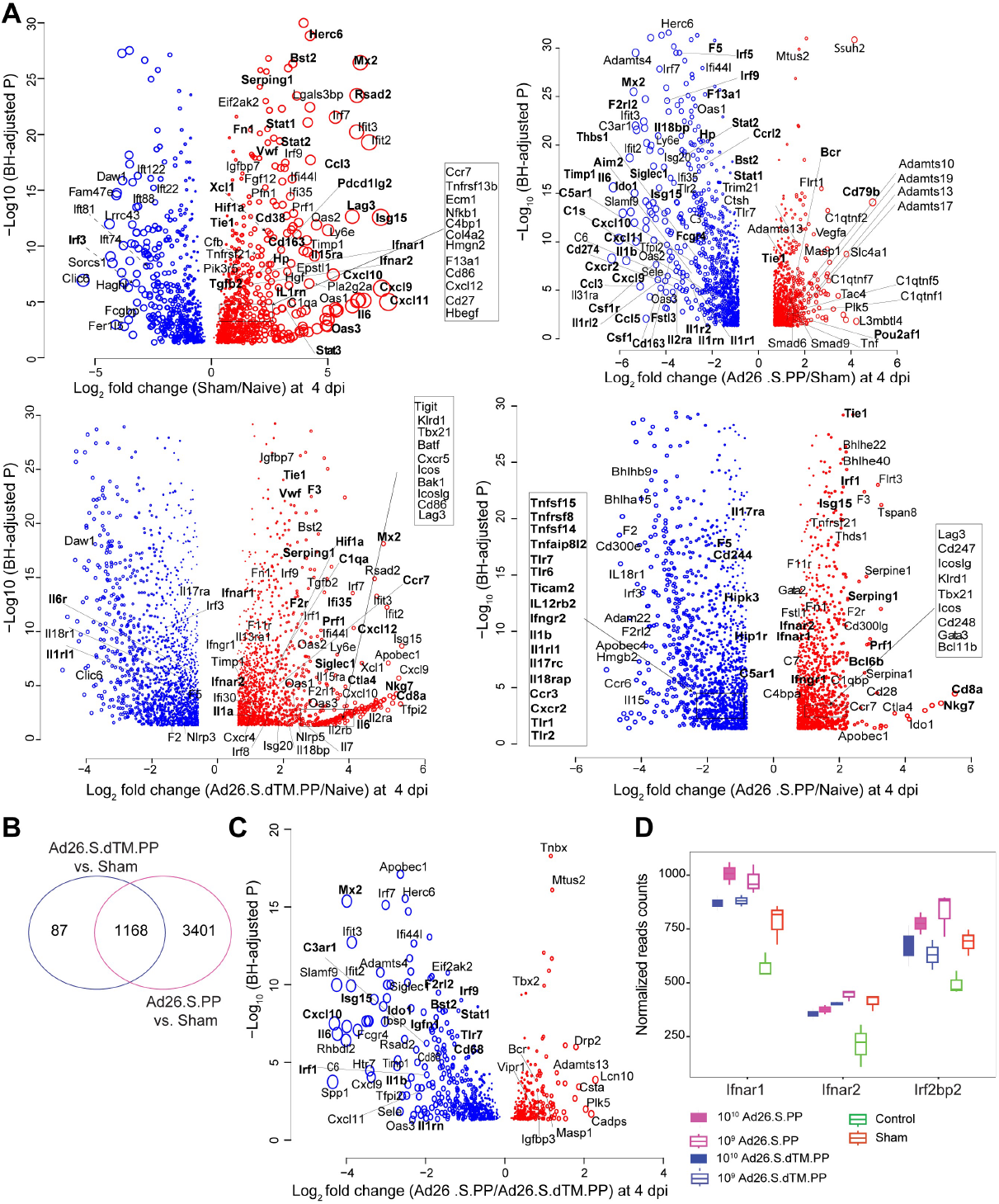
Transcriptomic changes in Ad26 vaccinated hamsters compared to sham unvaccinated and naïve control hamsters. **(A)** Scatter plots of differentially expressed genes in the lung of Ad26.S.PP, Ad26.SdTM.PP vaccinated and sham-unvaccinated hamsters compared to naive hamsters. Shown are genes that display a significant log2-transformed fold change. Coloration and point size indicate log2-transformed fold changes, red for upregulated genes and bleu for down regulated genes. The X-axis shows the log 2 transformed fold change expression. The Y axis, shows the -log10 adjusted p-values, of genes at 4 dpi. Adjusted P-values (<0.05) were calculated by DEseq2 using Benjamini-Hochberg corrections of Wald test p-values. **(B)** Venn diagram of common and distinct DEGs in Ad26.S.PP (pink circle) and Ad26.SdTM.PP (blue circle) vaccinated hamsters compared to sham animals at dpi. **(C)** Scatter plot of DEGs upregulated (in red) or downregulated (in blue) in Ad26.S.PP compared to Ad26.SdTM.PP at 4dpi. The X-axis shows the log 2 transformed fold change expression. The Y axis, shows the -log10 adjusted p-values, of genes at 4 dpi. Adjusted P-values (<0.05) were calculated by DEseq2 using Benjamini-Hochberg corrections of Wald test p-values. **(D)** Boxplot representation of the normalized reads counts of the interferon receptors 1 and 2 (Ifnar1 and Ifnar2) and interferon Regulatory Factor 2-Binding Protein 2 in vaccinated, sham-unvaccinated compared to naïve hamsters. Vaccination with two different doses were shown in full boxplots for the high dose and in empty boxplots for the low dose.

**S3 Fig.**
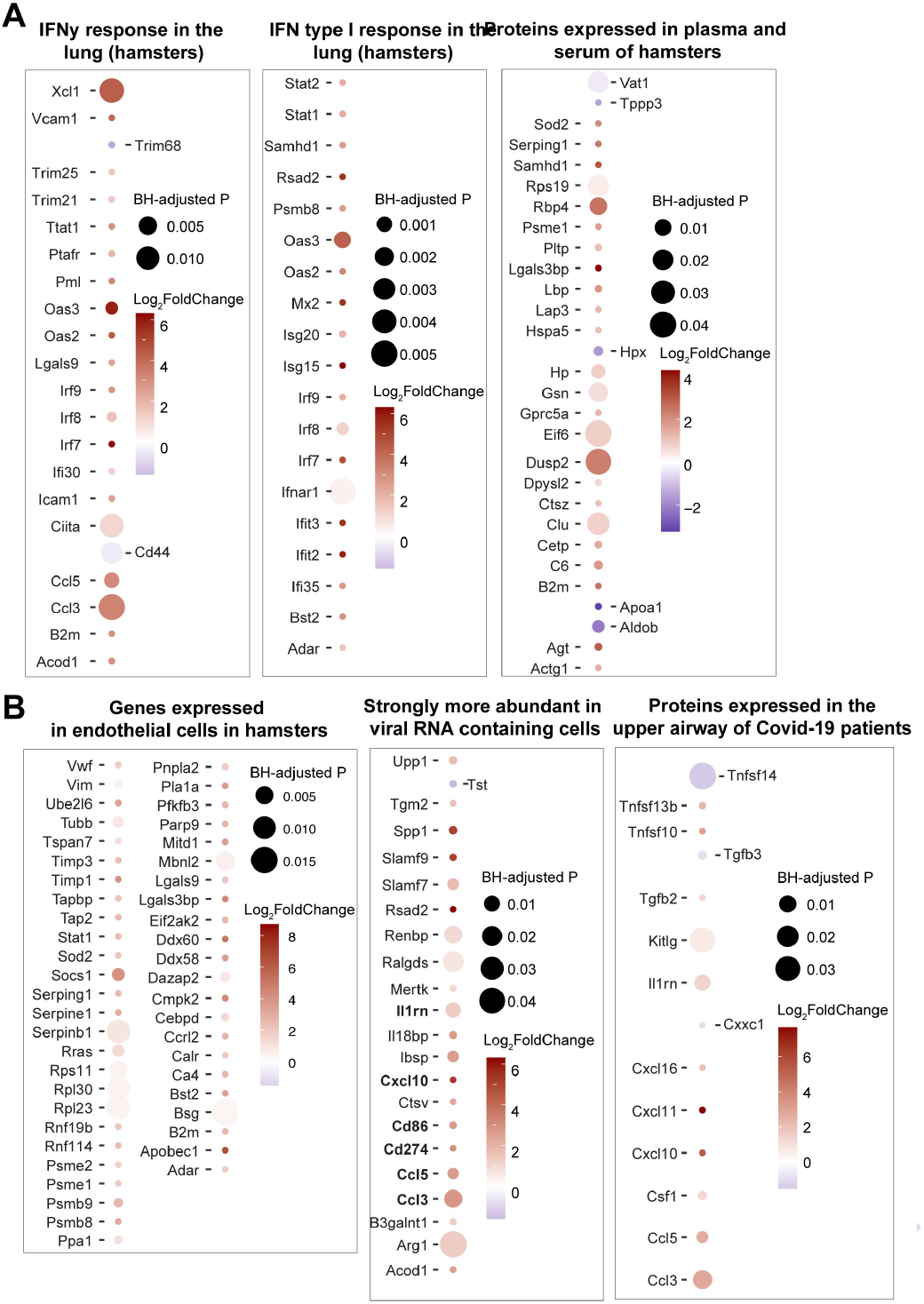
Our data recapitulate published omics analysis in hamsters infected with SARS-CoV-2, and in COVID-19 patients. **(A-B**) Dot plots of differentially expressed genes increased in infected hamsters at 4 dpi that are upregulated (red) or downregulated (blue) in COVID-19 infected patients, or in the lung of hamsters infected with SARS-CoV-2. Shown are genes that are increased (red gradient) or decreased (blue gradient) at 4 dpi in sham compared to naïve control animals, that were reported in a published SARS-Cov-2 hamsters study (6). Coloration and point size indicate log2-transformed fold changes and p-values, respectively, of genes at 4 dpi time point relative to control groups (naïve). Adjusted p-values were calculated by DEseq2 using Benjamini-Hochberg corrections of Wald test p-values.

**S4 Fig.**
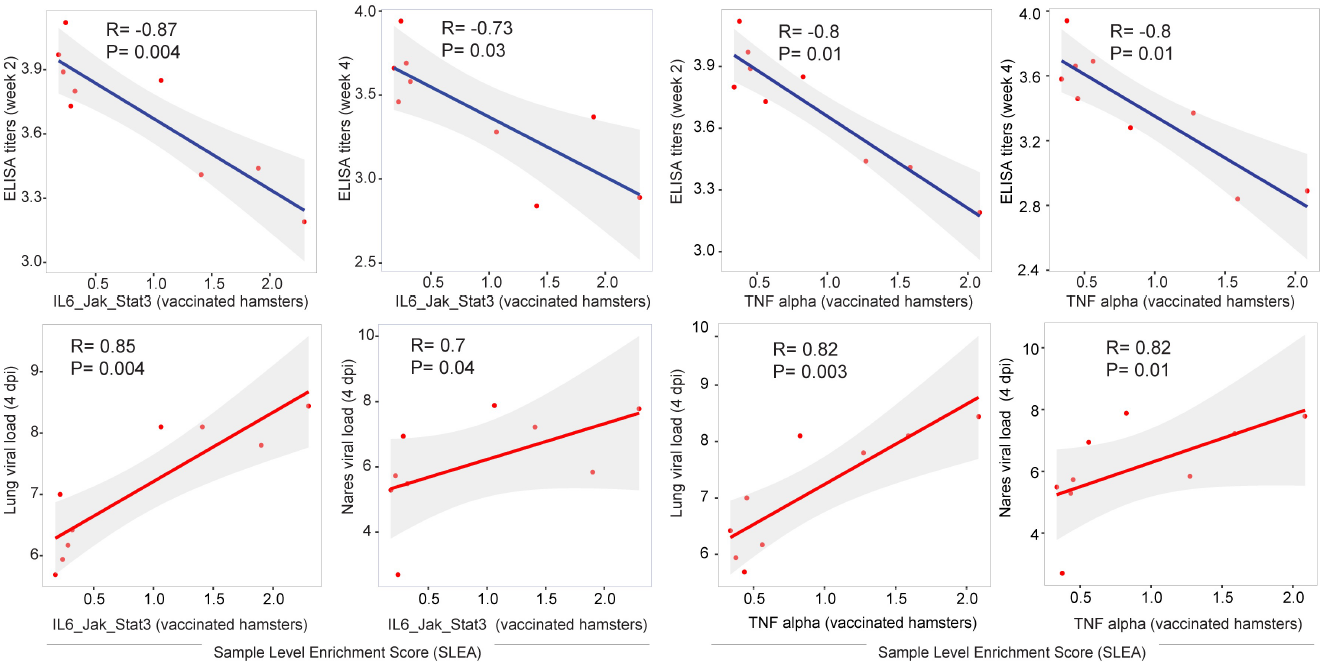
Proinflammatory pathways correlated positively with viral loads at 4 dpi in vaccinated hamsters. Scatter plot of the SLEA score of IL6_JAK_STAT3 and TNF alpha pathways in vaccinated animals at 4 dpi as a function of the ELISA binding titers, neutralizing antibody titers and viral load. The x-axis represents the samples level enrichment score of each pathway (SLEA) and the y-axis shows immune responses elicited by Ad26 in vaccinated hamsters. The average expression of the genes within each pathway was calculated using the SLEA z-score method. A linear regression model (blue or red line), and its 95% confidence interval (gray zone), was fit between SLEA z-score, viral load and the different antibody responses. A Spearman correlation and a t test were performed to assess the significance of the correlation between pathways SLEA scores and each response. Each red dot correspond to an individual animal. Positive correlations were shown in red and negative correlation were shown in blue.

**S5 Fig.**
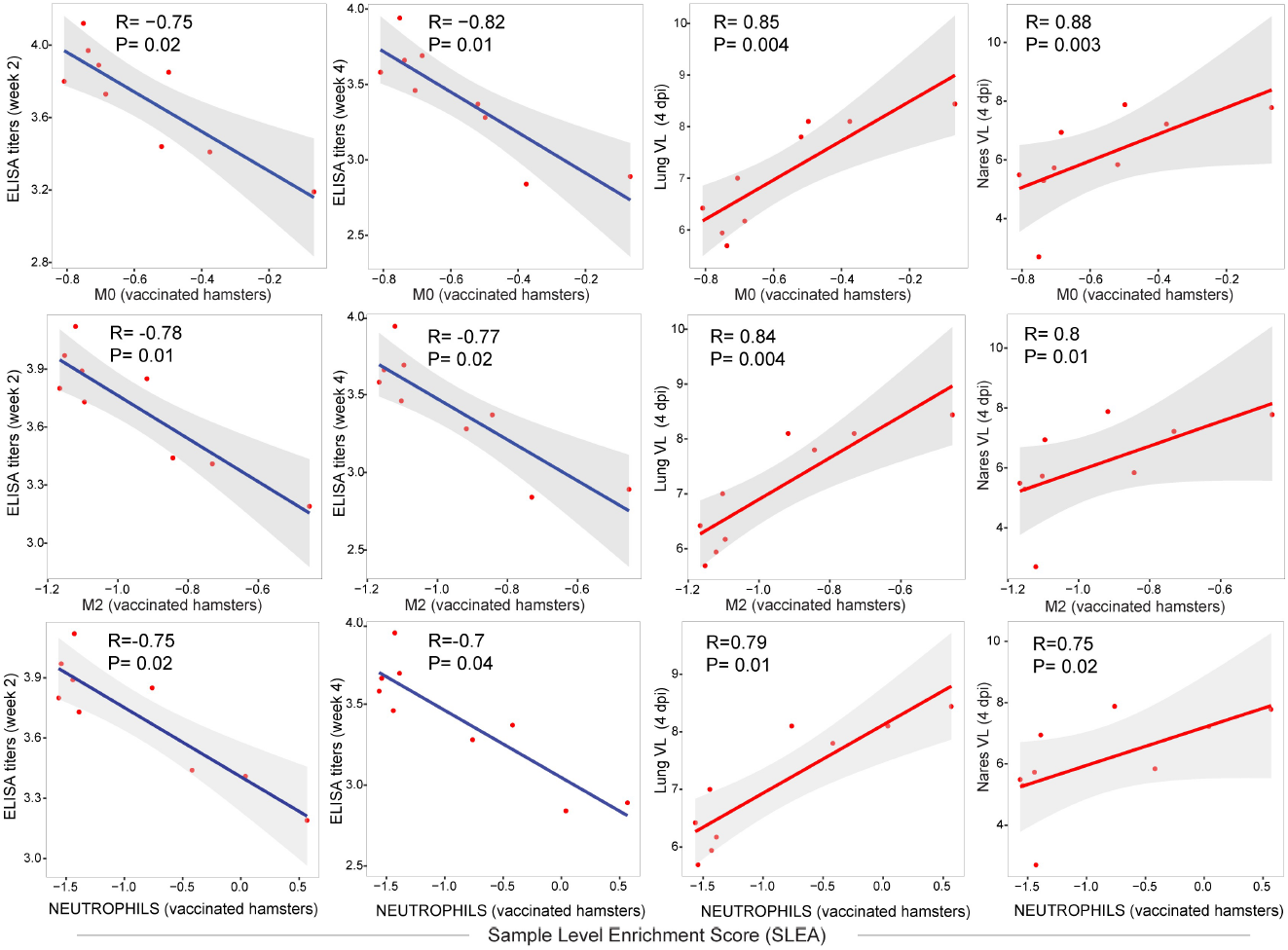
Correlation of macrophages and neutrophils signatures with binding and neutralizing titers and viral loads in vaccinated hamsters. Scatter plot of the SLEA score of macrophages pathways in vaccinated animals at 4 dpi as a function of the ELISA binding titers, neutralizing antibody titers and viral load. The x-axis represents the samples level enrichment score of each pathway (SLEA) and the y-axis shows immune responses elicited by Ad26 in vaccinated hamsters. P-value and the Spearman correlation coefficient were shown for each plot. Each red dot correspond to an individual animal. Positive correlations were shown in red and negative correlation were shown in blue.

**S6 Fig.**
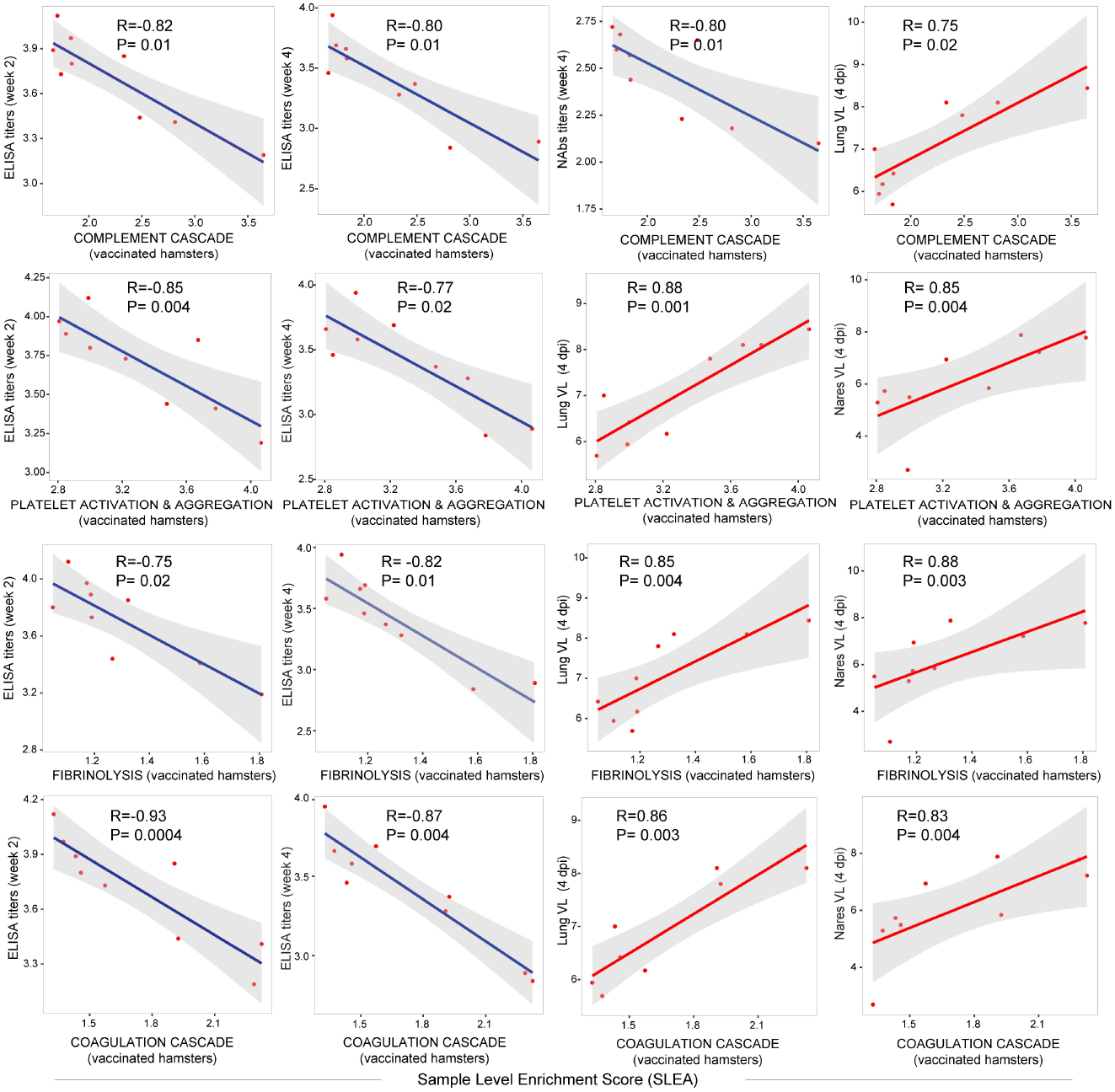
Correlation of thrombosis associated signatures with binding and neutralizing titers and viral loads in vaccinated hamsters. Scatter plot of the SLEA score of thrombosis associated pathways in vaccinated animals at 4 dpi as a function of the ELISA binding titers, neutralizing antibody titers and viral load. The x-axis represents the samples level enrichment score of each pathway (SLEA) and the y-axis shows immune responses elicited by Ad26 in vaccinated hamsters. P-value and the Spearman correlation coefficient were shown for each plot. Each red dot correspond to an individual animal. Positive correlations were shown in red and negative correlation were shown in blue

**S7 Fig.**
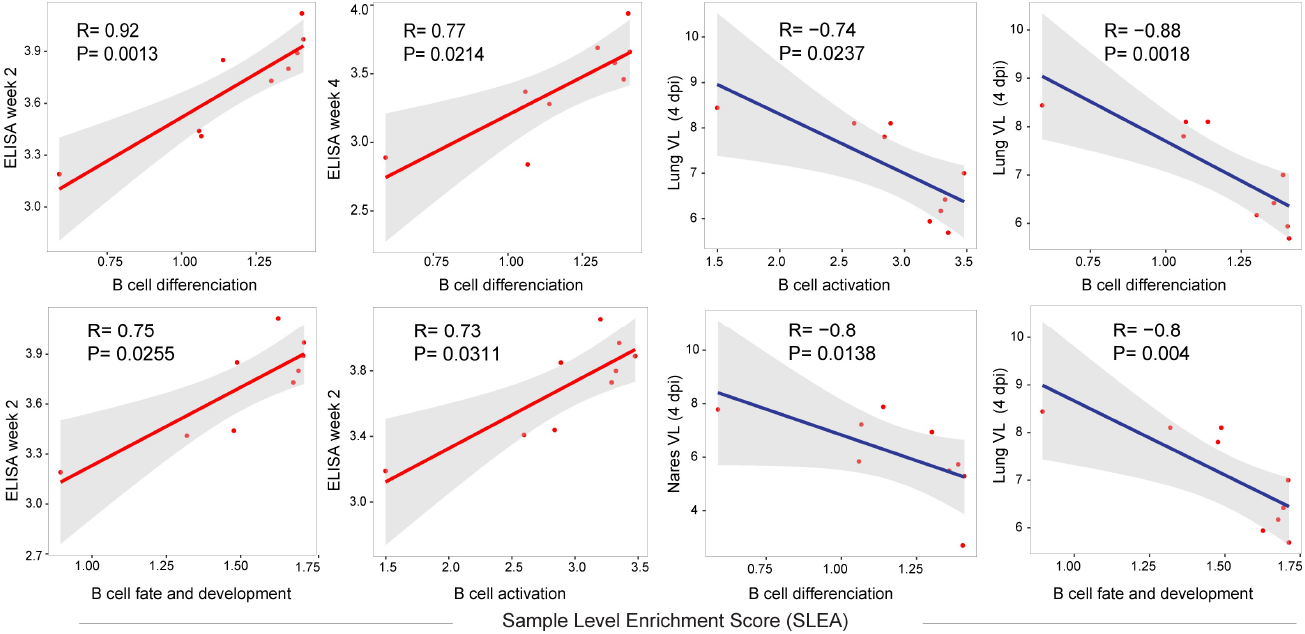
Correlation of B cell signatures with binding and neutralizing titers and viral loads in vaccinated hamsters. Scatter plot of the SLEA score of pathways of B cell activation, differentiation and B cell fate in vaccinated animals at 4 dpi as a function of the ELISA binding titers, neutralizing antibody titers and viral load. The average expression of the genes within each pathway was calculated using the SLEA z-score method. A linear regression model (blue or red line), and its 95% confidence interval (gray zone), was fit between SLEA z-score, viral load and the different antibody responses. A Spearman correlation and a t-test were performed to assess the significance of the correlation between pathways SLEA scores and each response. The x-axis represents the sample level enrichment score of each pathway (SLEA) and the y-axis shows immune responses elicited by Ad26 in vaccinated hamsters. Positive correlations were shown in red and negative correlations were shown in blue.

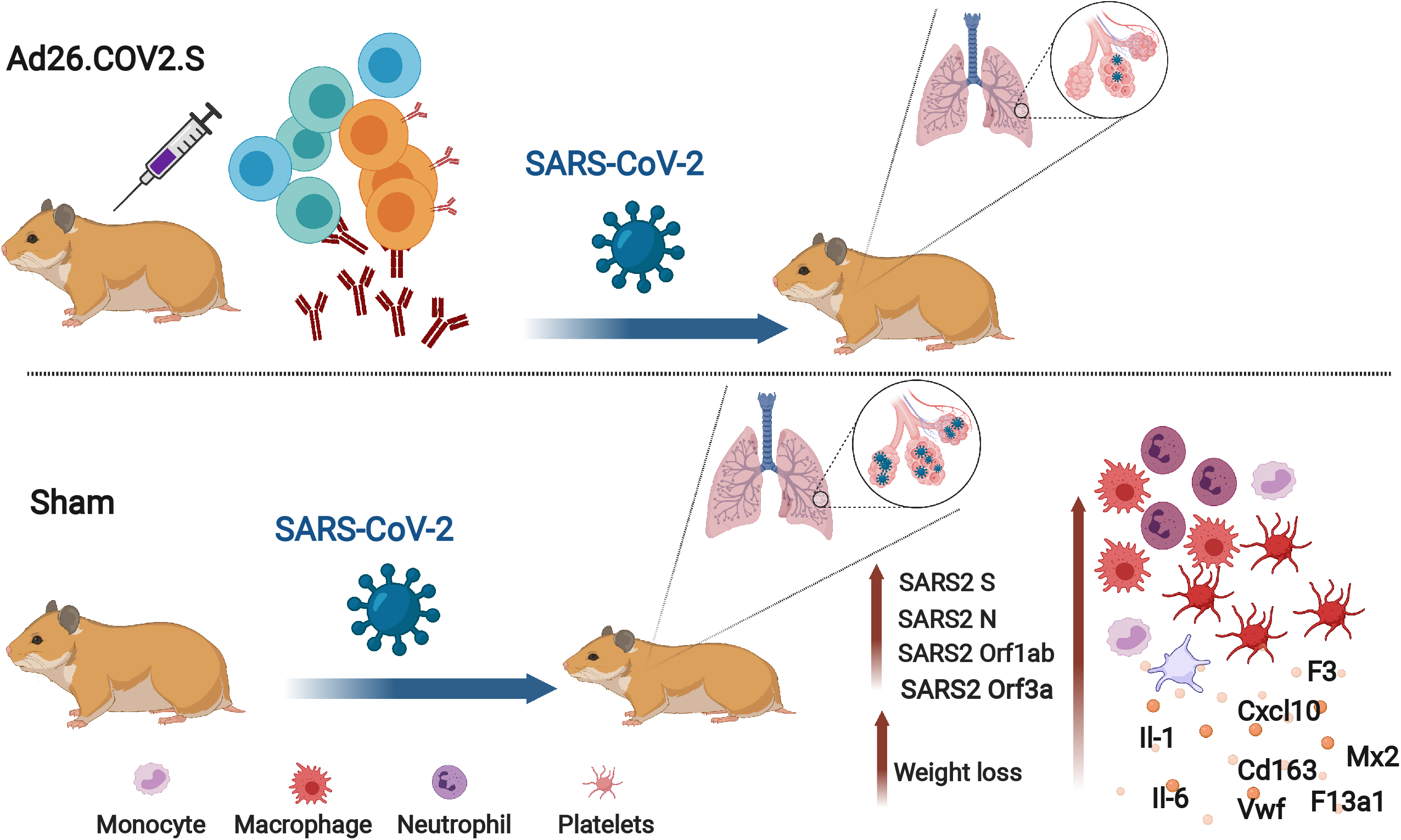

## Notes

### Competing Interest Statement

D.H.B. is a co-inventor on COVID-19 vaccine related patents.

